# Comparative analysis of root morphology in several spinach (Spinacia oleracea) varieties: Field vs Hydroponic growth systems

**DOI:** 10.64898/2026.04.07.717006

**Authors:** Deniz Camli-Saunders, Ava K. Russell, Camilo Villouta

## Abstract

Spinach (*Spinacia oleraceae*) is a principal vegetable crop commercially grown in Controlled Environment Agriculture (CEA). Recent research suggests that root morphological and architectural differences among crop species influence yield, resource use efficiency, and environmental stress tolerance. These root traits may be exploited to increase yield, promote efficient nutrient use, and mitigate environmental stressors. This study measured differences between various spinach cultivars in CEA systems to reveal morphological and anatomical variation. We grew three spinach cultivars with different reported growing rates (‘Income, ‘Darkside’, and ‘El-Majestic’) under NFT hydroponic and substrate-based systems in a controlled greenhouse environment over 45 days with destructive harvests at days 15, 30, and 45. Supplemental light (250 µmol/m^2^/s) with 12-hour photoperiod and periodic fertigation was used. Harvests included the collection of leaf and root biomass, and scanning of root systems in WinRhizo software, measuring ten variables. On day 45, root cross-sections from orders 1-5 were embedded in JB-4 resin, sectioned, stained, and analyzed for diameter, vasculature, and rhizodermis characteristics. Results indicate that in spinach, differences in root system morphology are linked to cultivation systems over cultivar identity. Vascular and root anatomical alterations are minor compared to morphological differences in response to the cultivation system. Hydroponic-style growth systems are associated with the proliferation of fine-root ideotypes compared with substrate-based conditions. Such findings affirm previous studies, which suggest plastic root morphology in response to growth systems, and may be used to help create more resilient, resource-efficient cultivars.

**Highlights:**

1. In spinach, root system morphology differences are linked to cultivation systems.
2. Root vascular and anatomical alterations are minor in response to cultivation system.
3. Hydroponic growth systems are linked to fine-root ideotype proliferation in spinach.
4. Fine-root ideotype proliferation may be a breeding target for CEA spinach.

## 1. Introduction

Controlled Environment Agriculture (CEA) is a pillar of sustainable agriculture initiatives around the world (Sharma et al., 2024; Fussy and Papenbrock, 2022; Engler and Krarti, 2021). CEA allows growers to curate ideal conditions for a range of crops, significantly increasing yield and harvest frequency while combating unpredictable climatic trends (Cowan et al., 2022; Benke and Tomkins, 2017). Such conditions include ideal temperature ranges, satisfactory supplemental lighting (often via Light Emitting Diodes), and highly oxygenated root zones (Paponov and Paponov, 2025; Vega et al., 2025; van der Ent et al., 2024). Spinach, *Spinacia oleracea* L., is among the most commercially significant leafy vegetable crops in CEA production owing to its short growth cycle, versatility across production systems, and high nutritional and market value (Tai et al., 2020; Qin et al., 2017; Citak and Sonmez, 2009). However, there is a limited understanding of spinach root systems in CEA. It remains unclear how the cultivation system (nutrient film technique, NFT, versus substrate-based) and cultivar growth habit relate to root order distribution, morphology, and vascular anatomy. Our study aims to bridge these gaps by characterizing root morphological and anatomical traits of three spinach cultivars (with fast, medium, and slow growth rates) across NFT and substrate-based CEA conditions. Based on prior studies, we believe that the growth environment will outweigh cultivar-based differences, meaning that plants grown in hydroponic production systems will have finer, more highly branched, and longer root systems.

Root system architecture plays a central role in plant performance through its regulation of resource acquisition, transport, and allocation (Zhou et al., 2022). Prior literature has established the significance of root architecture in CEA systems, though mainly in lettuce and herb crops. Root systems can be broadly categorized by function: absorptive roots, with wherein roots take up resources and nutrients, and transportive roots, wherein roots facilitate axial movement of these resources (Li et al., 2023 McCormack et al., 2015). These functional distinctions are associated with specific morphological and anatomical traits including root diameter, surface area, and degree of lateral branching. Exploration can be defined in part by the incidence of root tips and forks. Root tips act as sensory organs, allowing the flow of tactile information about environmental makeup and stressors (Yokawa & Baluska, 2018). Forks, the split of a root into a separate branch represents a decision in regards to stimuli: whether environmental or chemical, that drove the root system to continue an exploration to a certain area (Danjon & Reubens, 2008). A main method of organizing root systems is through root order classification. This study adapted the methods set out in Atkinson et al., (2014), and Meng et al., (2018), wherein lateral roots increased in order from proximal to distal (Fig. 1). Finer roots, such as 3^rd^, 4^th^, and 5^th^, order roots, help increase overall surface area, and act as nutrient absorption points, thereby enhancing nutrient foraging capabilities (Li et al., 2023, Chen et al., 2013). Thicker roots, like first and second order roots, have greater conductance and support axial transport capacity via larger stele area and xylem development (Balliu et al., 2021, Gu et al., 2014). The composition and distribution of these orders among various cultivars together represent a primary determinant of growth capacity, resource use efficiency, and stress tolerance. Such compositions and distributions, specifically their conductance structures, have yet to be investigated or compared in CEA-grown spinach among substrate-based vs hydroponic conditions.

**Figure 1:**
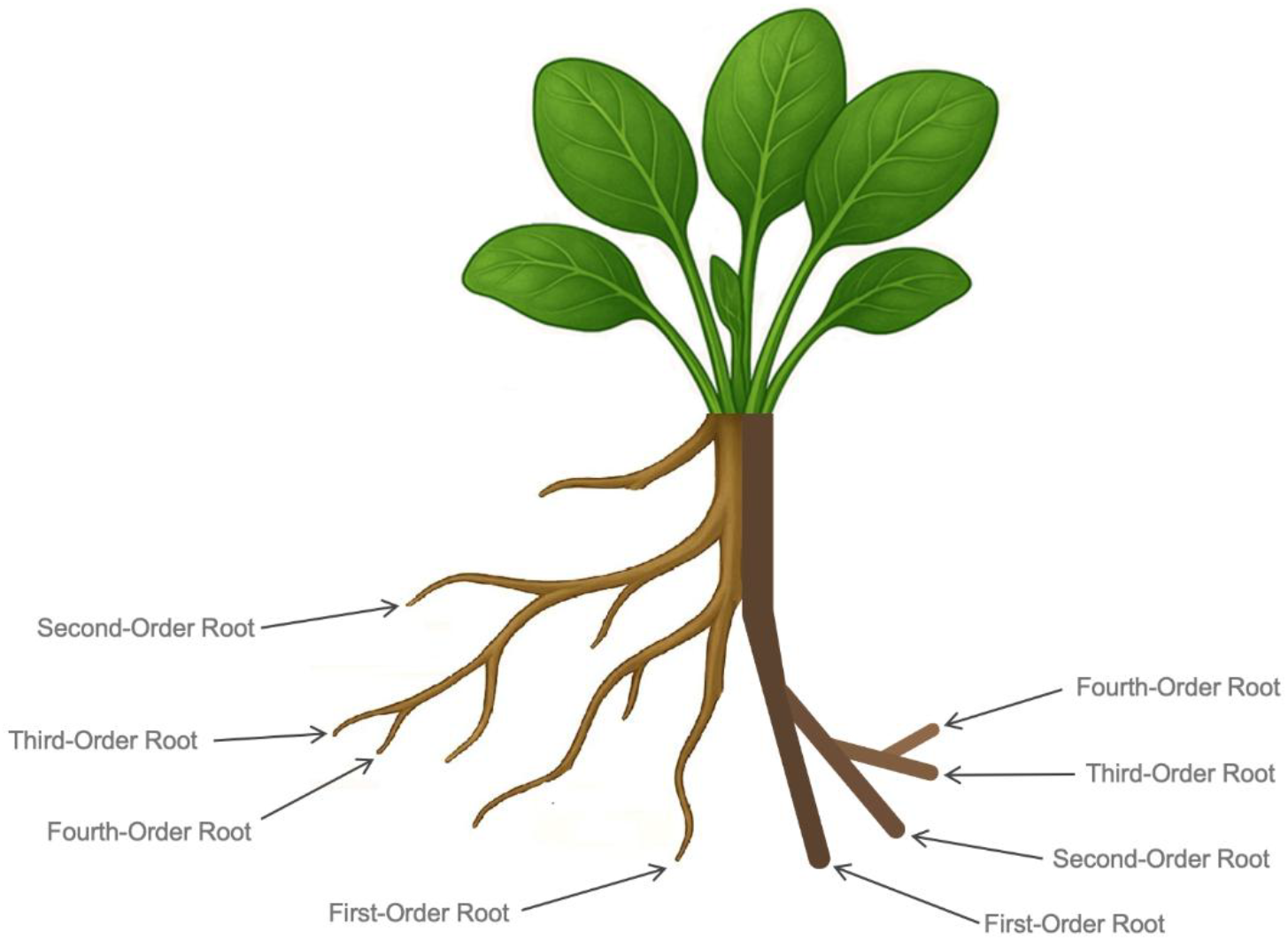
Schematic representation of root order classification applied to spinach (*Spinacia oleracea*) Schematic root order distribution of spinach. Root order was determined through the adaptation of the protocols established by Atkinson et al. (2014) and Meng et al. (2018), wherein the most proximal root represents the first order and increases toward the most distal. As a dicot, spinach roots have a main taproot with several lateral roots branching from it. The main taproot is considered first order, and with each successive branching, root order increases. A lateral root branching from the main root would be second order, and a root branching from that would be considered third order. First- and second-order roots are noticeably thicker than third-, fourth-, and fifth-order roots. As root order increases, root functionality shifts from transport-based to absorption-based.

Efforts to describe common root architectures have led to the development of root “frameworks” which encapsulate various morphologies, strategies, and distributions. These have been limited mostly to field grown crops. One framework developed in field conditions is the Subsoil Foraging Ideotype (SFI), colloquially known as “Steep, Deep, and Cheap.” SFI is characterized by an efficient water and nutrient acquisition network anchored by a strong primary root, high lateral development, and a balanced absorptive to transportive root ratio (Lynch, 2013). The development of SFI traits has been linked to environmental stress tolerance, improved nutrient uptake, and increased overall productivity (Koevoets et al., 2016). These root “frameworks” have yet to be adapted for hydroponic conditions, with descriptions remaining vague. However, such traits, including improved nutrient uptake and overall productivity may still be advantageous in hydroponic growth environments.

Spinach root anatomy directly influences hydraulic conductance and transportive capacity, with key traits including xylem vessel diameter, stele area, xylem number, and overall vascular cross-sectional area (Peng et al., 2024; Chen et al., 2013). The understanding of spinach crop production in CEA systems is in its initial stages. Several important areas of study, including root architecture, order differentiation, and functional traits, have yet to be explored and may serve to optimize and improve harvest turnover and resource efficiency. Novel studies of other crops, including tomato, have revealed differences in xylem diameter, stele size, xylem number, and overall conductance, especially under differing growth conditions (Ayarna et al., 2020). In cereal crops, studies have been primarily focused on anatomical responses under drought or flooding (Ranjan et al., 2022; Fonta et al., 2022; Bacanamwo & Purcell, 1999). Furthermore, these studies are often limited in their use of microscopic cross-sectioning analysis. Of those that do employ the technique, they are often focused on rice or wheat (Fonta et al., 2022; Ouyang et al., 2020). Under drought conditions, rice shows comparatively limited anatomical plasticity under water deficit, whereas wheat exhibits xylem-level adaptation in response to varying soil moisture (Ouyang et al., 2020). In Sorghum, anatomical plasticity in response to drought was similarly limited (Guha et al., 2018). Among vegetable crops, such as lettuce, anatomical plasticity appears to be only modest between hydroponic and soil conditions, whereas cereal crops such as wheat have exhibited major anatomical plasticity (Lei & Engeseth, 2021; Kadam et al., 2017). Differences appear to be highly context dependent, or rooted in genetics, in contrast to morphology which shows clear-cut differentiation between systems (Schneider et al., 2020). A baseline of plasticity in spinach may be established by examining the anatomy of varying cultivars under multiple growing conditions.

Environmental growth conditions play a major role in root morphological and anatomical differentiation and development. Consistent nutrient availability, access to water, and an oxygen rich environment like that of a hydroponics system, create a cascade of architectural adjustment, including the thinning of root cortexes, branching of lateral roots, and overall root system lengthening. The high, homogeneous nutrient availability, together with elevated dissolved oxygen (DO) levels, enables pericycle cell activation, signaling the root system to undergo lateral growth (Striker, 2023; Winkel et al., 2013). To accommodate this lateral growth, cells in the cortex may expand, thereby increasing the overall root diameter and facilitating solute transport. Resultantly, hydroponically grown crops, particularly vegetable crops, exhibit increased lateral root growth (visible by the 3-5^th^ order roots) as compared to soil-grown counterparts (Sasse et al., 2020). Such high DO conditions may also reduce or eliminate the formation of aerenchyma in cortical spaces, thereby “thinning” the overall root cortex (Pan et al., 2021). Moreover, the virtual elimination of mechanical resistance in hydroponic systems in comparison to soil-grown counterparts further thins the root cortex (Nitu et al., 2024; Scott and Villouta, 2026). The aforementioned cascade of alterations may result in increased root surface area which allows for higher overall rates of absorption and transportation (Nitu et al, 2024; Que et al., 2018). Higher transportation may be achieved through adjustments in the transportive roots caliber, increasing the average stele diameter to accommodate larger transport conduits (Balliu et al., 2021; Guo et al., 2008) From this, it becomes clear that architectural specialization clearly occurs in response to the growth environment, and that these root systems, particularly in vegetable crops, are highly plastic.

This sequence of morphological changes in response to the environmental conditions can be predicted with further understanding and investigation. Better determination of vegetative growth productivity, and root architecture could reveal the variables that determine important values such as: yield, harvest turnover, and resource efficiency (Lombardi et al., 2021). Our study aims to shrink these unknowns by establishing root morphological characteristics and determining differences in inner anatomy under NFT vs substrate-based conditions across three variable speed spinach cultivars. In accordance with prior literature, we believe root morphological differences based upon growth environment will outweigh cultivar-based growth differences, meaning that plants grown in hydroponic production systems exhibit higher plasticity in the form of finer, longer root systems.

## 2. Materials and Methods

### 2.1 Plant Material and Experimental Structure

Three commercially relevant cultivars of spinach. ‘Income’ was a fast-growing cultivar, ‘Darkside’ was an average/middle growth speed cultivar, and ‘El-Majestic’ (‘Majestic’) was a slow-growing, heat tolerant cultivar.All plants were grown in a walk-in greenhouse at the University of Rhode Island (URI’s Greenhouse Complex, Kingston, RI) for this study. This experiment followed a completely randomized design (CRD), with two cultivation systems as the primary treatment factor.

The plants were grown under environmentally controlled conditions of 12-hour photoperiod (250 µmol/m^2^/s), at an average temperature of 25 °C in either a modified NFT hydroponic system (CropKing, Ohio USA) or in one-gallon plastic pots with Pro-Mix Mycorrhizae soil mix (Premier Tech, Rivière-du-Loup, QC, Canada). Hydroponic fertigation was via a sump tank containing equal parts Jack’s Part A (5-12-26) and Part B (15-0-0) with pH 6.2-6.7 and EC 1.6-1.9.) Soil-based trials were fertigated once daily with 300mL of the same Jack’s Part A & B solution with the same pH and EC ranges as hydroponic systems. A total of 10 plants were grown per cultivar per harvest date per system, randomly distributed across 6 NFT systems, and harvested destructively at three time points (days 15, 30, and 45 post-sowing).

Prior to planting, the seeds were removed from cold storage (4 °C) in order to break dormancy and imbibed in a 10% solution of hydrogen peroxide (H_2_O_2_) for a period of two hours, followed by a 4-hour soak in De-Ionized (DI) water. For the hydroponic trials, seeds were then planted into lightweight expanded clay aggregate (LECA) media inside a 2-inch net pot. The net pots were then placed into NFT gutters. For the soil trials, seeds were potted into one-gallon pots.

### 2.2 Plant Processing and Root Imaging

Destructive harvests were conducted at 15, 30, and 45 days after sowing (DAS) with ten plant per cultivar per system collected at each time point. For hydroponic systems, root systems were carefully removed from the net pots and LECA medium without washing and proceeded directly to the weight collection stage. Soil trials underwent a brief wash protocol to remove soil from the root systems and ensure high-quality image collection. The wash protocol included three stages. First, a manual soil-removal stage in which soil was raked out of the root system by hand. Second, roots were rinsed in a container of tap water for 1 minute. Third, a second rinse in a fresh container of tap water for another 1 minute, with light manual agitation to remove clinging soil particles. At that point, roots were allowed to dry for 5 minutes before fresh weights were recorded on a precision balance (Entris® II Essential Line BCE6202I; Sartorius AG, Göttingen, Germany; 6,200 g capacity; 0.01 g readability). Weight collection included fresh weights of both leaf and root material, as well as dry weights collected after 24 hours in a mechanical convection oven (Heratherm OMH180-S; Thermo Fisher Scientific, Waltham, MA; 6.3 cu ft, 120 V). After fresh weight collection from both growth systems, roots were placed on WinRhizo trays and separated with small spatulas and forceps to preserve root structure. Larger root systems were cut with razor blades to ensure accurate visualization of roots. Upon completion of root spreading, the trays were scanned in an Epson Expression 12000XL scanner with the WinRhizo Pro analysis program (Regent Instruments, Quebec, Canada). The analysis function was used to collect measurements, including total root system length, average and measured diameters, volume, surface area, tips, forks, crossings, and root diameter classification. Root systems that required multiple sequential scans were merged using the “XLRhizo” aggregation module.

Data was exported to Excel for processing and grouping. Later, it was uploaded to the data analysis program “R” (version 4.5.0;R Core Team, Vienna, Austria) and analysis was performed using R packages: “ggplot2,” “dplyr,” and “tidyverse” among others. Two- and three-way Analysis of Variance (ANOVA) was performed, along with Tukey’s test for instances where the main effect was statistically significant. Further analysis of several two and three-way interactions was conducted to compare main effects in differing growth systems (i.e., Hydroponic vs Soil-based). Plots were created to visualize specific measurements/traits per harvest day and per growth system.

### 2.3 Histological Preparation and Microscopic Analysis

On harvest day forty-five, four plants per cultivar were randomly selected for histological analysis. Root segments were collected from each plant representing orders 1 through 5 following the order classification framework of Atkinson et al. (2014), wherein first-order roots are the most proximal to the root–shoot junction, and order number increases distally (Fig. 1). Segments were placed in 20 mL scintillation vials with 1.5 mL of 4% Paraformaldehyde in PBS (Thermo Fisher Scientific, Waltham, MA, USA) solution. Samples underwent three treatments in a vacuum desiccator connected to a diaphragm vacuum pump (Model GM-0.5A, Huanyu, Shenzhen, China) for 2, 5, and 10 minutes, respectively, at ∼10 psi to ensure proper infiltration. After vacuum treatments, samples were placed on an orbital shaker and stored overnight at 4 °C.

On the following day, the samples were removed from the refrigerator and underwent an ethanol titration protocol beginning with PBS wash (x3) and ethanol concentrations of 10, 20, 30, 50, and 70%, respectively for a period of one hour each. For full protocol, see supplementary materials.

Root segments were prepared for infiltration with JB-4 glycol methacrylate resin (Sigma-Aldrich, St. Louis, MO, USA). Samples were further dehydrated with one-hour increments of 85%, then 95% ethanol (x3) before infiltration began with 3:1 solution of 95% EtOH-MonoA solution. Infiltration continued over four days with daily replacements of solutions with increasing concentrations of Mono A (Day 1 3:1, Day 2 1:1, Day 3 1:3 EtOH:MonoA) before concluding with 100% Mono A solution. The solution of Mono A was replaced weekly for a period of three weeks before the embedding/polymerization steps could begin.

Embedding was performed with fresh Mono A solution, as described in Sigma-Aldrich technical bulletin. Samples were placed in molds, and polymerization occurred under anaerobic conditions as specified. The polymerized samples were then mounted to acrylic stubs and labeled. Samples were sectioned at 1.5 µm thickness in a humidity-controlled room set at 70% RH with a rotary microtome (Model HM355S, Epredia, Leica Biosystems, Germany) with fractured (triangular) glass knives. Cut sections were placed on a slide containing DI water drops to relax and expand, and dried overnight at 45 °C on a slide warmer. Slides were stained using a 0.1% Toluidine Blue w/v solution. Toluidine Blue was pipetted onto slides for 45 seconds, then rinsed with DI water and allowed to air-dry. After staining and drying, coverslips were mounted with Permount (Fisher Scientific, Hampton, NH, USA) in a fume hood and allowed to dry for a period of at least twenty-four hours. Slides were analyzed using a Zeiss Imager A2 Microscope, with images captured and exported to Image J software (Schindelin at al., 2012). Images were analyzed for the following traits: root diameter, stele diameter, number and average diameter of xylem. Data was exported to Excel and underwent analysis.

### 2.4 Environmental Conditions Data Collection

Environmental conditions were continuously recorded throughout the 45-day experimental period. Temperature was measured using a Hobo Pendant (Model MX, Licor, Nebraska USA). Sensors were positioned at canopy level within multiple growing systems. Air temperature, sensors were housed under radiation shields and suspended from gutter structure. Data was collected at 5-minutes intervals for the duration of the experiment. Water reservoir dissolved oxygen (DO) content was measured in the NFT sump tank using a portable dissolved oxygen meter (Model HI98193; Hanna Instruments, Woonsocket, RI, USA). Three independent measurements were taken per tank under full-flow operating conditions and averaged.

## 3. Results

### 3.1 Morphological Analysis

Ten variables were analyzed to assess their statistical contributions to morphological differences in this experiment. Of particular note were the following: average root length, average root surface area, average root diameter, average fresh leaf weight, and average dry root weight. All analyses indicated harvest based main effect differences due to the temporal nature of the periodic harvest. Total root length had multiple main and two-way interactions including by System (P<.001) and Harvest:System (P<.001). Hydroponically grown plants had consistently longer total root lengths than their soil system counterpart (Fig. 2D). Harvest one had cultivar ‘Darkside’ as the longest, harvest two was ‘Majestic’, and harvest 3 was ‘Income’. Similarly, average root surface area had significant main effect for system (P<.001), nearly significant main effect for cultivar (P<.066) and significant interactive effects for Harvest:Cultivar (P<.006), Harvest:System (P<.001), Cultivar:System (P<.013) and 3 way interaction P<.001. Hydroponically-grown plants had higher root surface areas than their soil-grown counterparts across all cultivars (Supplementary Fig. 2D) and harvests save harvest three where cultivar ‘Income’ had larger root surface area for soil-grown plants. Significant differences were different in all main and interactive effects for average root diameter across all three harvests. Post-hoc analysis indicated that hydroponically grown plants had average root diameters smaller than their soil counterparts, and slow-growing cultivar ‘Majestic’ consistently had the largest root diameter of all cultivars across all harvests (Fig. 2C). These differences are clear in the interactive effects of Harvest:Cultivar (P<.001), Harvest:System (P<.001), Cultivar:System (P<.001) as well as three-way interaction (P<.0017). Such trends demonstrate that soil-grown plant roots have lesser development of 3rd, 4th, and 5th order roots as compared to hydroponically grown counterparts. Dry average root weight also demonstrated clear systemic differences (P<.001) apparent in Supplementary Figure 2B. Hydroponically grown plants consistently had increased dry root weights as opposed to soil-grown plants, across all cultivars and harvests with the exception of cultivar ‘Income’ in harvest three, wherein the soil dry root weight was larger than hydroponic weight. The aforementioned morphological differences translated into vegetative growth differences as well. Across all harvest and cultivars, hydroponically-grown plants had higher fresh leaf weights than soil-grown counterparts (P<.001)(Fig. 2A). System based main (P<.001) and three-way interaction, P< .019) were also significant.

**Figure 2:**
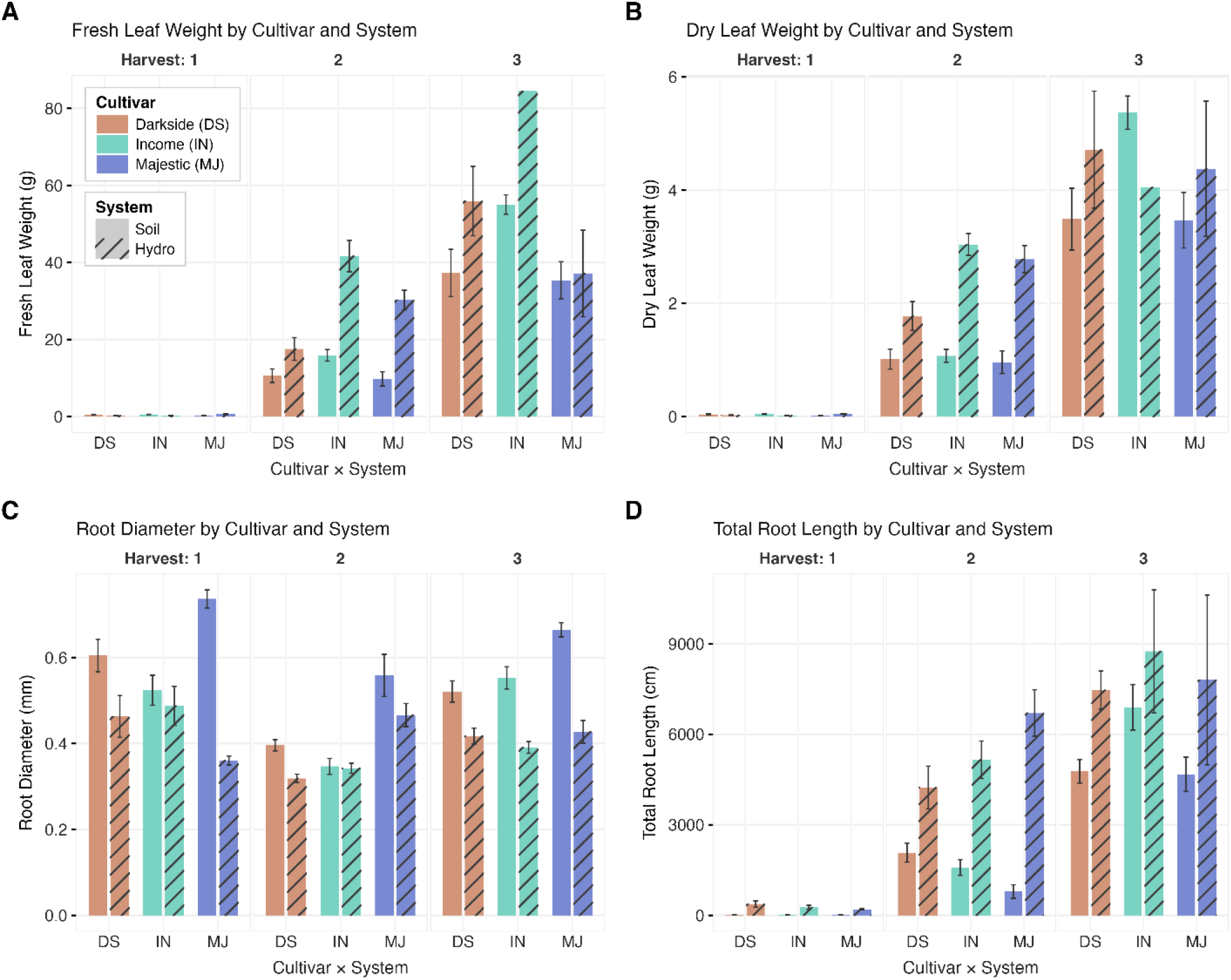
Morphological development of spinach (*Spinacia oleracea*) across three harvests under hydroponic and soil growing systems. Bar plots illustrating morphological root traits and aerial biomass across harvests. Each panel represents three cultivars — orange: “Darkside” (DS), teal: “Income” (IN), blue: “Majestic” (MJ), by growing system: solid = soil, striped = hydroponic, and by periodic harvest. Harvest 1 (day 15) is the leftmost section of each panel, harvest 2 (day 30) is the center, and harvest 3 (day 45) is the rightmost. Error bars represent ± 1 standard error of the mean (n = 10). Asterisks (*) indicate statistically significant differences between growing systems within the same cultivar. Data aggregation, statistical analysis, and figure generation were performed in R. Panel A: Fresh leaf weight immediately after harvest. Panel B: Dry leaf weight collected after 24 hours at 60°C in a drying oven. Panel C: Average root diameter (mm) of the entire root system per treatment and harvest, measured using a WinRhizo root scanner. Larger average root diameters are indicative of root systems dominated primarily by first- and second-order roots rather than third-, fourth-, and fifth-order roots. Panel D: Total root length (cm) per treatment and harvest, measured using the WinRhizo root scanner. Wide variability was observed at harvest 3; however, a consistent trend of hydroponically grown root systems being longer than their soil-grown counterparts was evident across harvests.

Other variables which also demonstrated significant differences were root volume (cultivar main effect P<.00012, Cultivar:System, P<.037) (Supplementary Fig. 2C), number of tips (cultivar main effect, P<.001, harvest:cultivar P<.001), number of forks (system main effect, P<.001, harvest:system, P<.0003) and number of crossings (system main effect, P<.001, harvest:system P<.001) (Supplementary Fig. 1B,1C). These latter two support the exploratory nature of root morphology in hydroponic systems.

### 3.2 Anatomical Analysis

Anatomical analysis through microscopic observation led to the collection of six separate variables of collection per root order. These were the number of xylem, average individual xylem area, average individual xylem diameter, root diameter, stele diameter, and a vascular tissues ratio (stele diameter divided by root diameter). Statistical analyses of these results revealed significant three-way interactions for the following variables: average xylem area (P<.0038), average xylem diameter (P<.0026), and near significance for stele diameter (P<.06). Due to the presence of numerous smaller proto-xylem in the first order roots, second order roots had the highest xylem area on average. Of all cultivars in second order roots, ‘Income’ had the largest xylem area. However, in orders 3-5, cultivar ‘Darkside’ had the largest xylem area on average (Supplementary Fig. 3B), followed by ‘Income’, and then cultivar ‘Majestic.’

While three-way interactions of xylem diameter were demonstrated to be significant, no clear trend emerged by system or order as to which was consistently larger. In order one, ‘Darkside’ soil roots had the largest xylem diameter, followed by ‘Income’ hydroponic, while in order two ‘Income’ hydroponic and soil roots were largest. ‘Majestic’ consistently had the smallest average xylem diameter in orders 1-4, while in order five these numbers evened out and ‘Majestic’ hydroponic roots barely became the largest.

For stele diameter, a trend emerged specifically in orders one, two, and five. In these orders, hydroponic roots had higher average stele diameters than the same cultivar in a soil system. In order three, this trend disappeared, and in order four reversed with soil stele diameter being slightly larger on average than hydroponically grown plant roots. There were several notable significant pairwise comparisons in measurements of stele diameter. Hydroponic first order roots of ‘Darkside’ had larger stele diameters than their soil counterparts. Hydroponic second order roots of ‘Majestic’ also had larger stele diameters than their soil counterparts. Pairwise comparisons in orders 3-5 did not reveal any similar differences.

Though not significant in pairwise or interaction comparisons, the number of xylem decreased with each order, ranging from 50-70 for first order roots of all cultivars, and falling to 3-8 by the fifth order of roots (Supplementary Fig. 3A). Root diameters between cultivars remained similar while decreasing in each order (Figs. 3 and 4). While not significant, ‘Majestic’ had the largest average root diameter across all three cultivars and five orders (Fig. 4). The average vascular tissue ratio had one clear trend emerge, ‘Income’ hydroponic roots had a higher makeup of their cells as vascular tissue than their soil counterparts across all orders (Fig. 4). This trend was also seen for ‘Darkside’ in first, third, and fifth order roots, and in ‘Majestic’ in first and second order roots.

**Figure 3:**
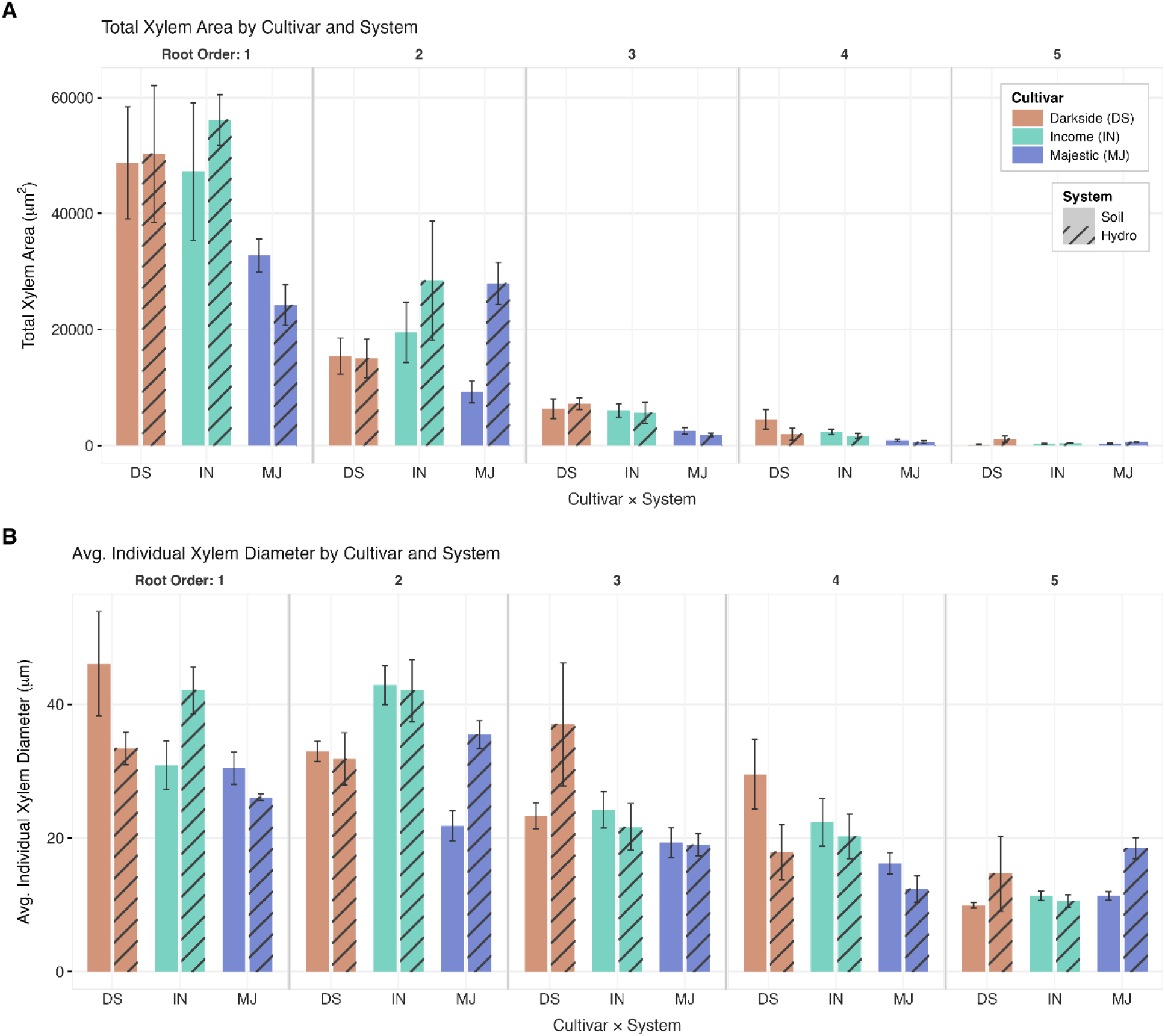
Xylem anatomy of spinach (*Spinacia oleracea*) roots across orders and growing systems. Analysis of root vascular anatomy across root orders and growing systems. Samples for each root order, cultivar, and growing system (n = 4) were fixed, dehydrated, embedded in JB4 resin, sectioned at 1.5 μm thickness, and stained with toluidine blue. Imaging was conducted using a Zeiss Imager A2, with image processing performed in FIJI and data analysis in R. Error bars represent ± 1 standard error of the mean. Panel A: Total xylem area (µm^2^) per treatment and root order (1–5). Cultivars are denoted by color: orange: “Darkside” (DS), teal: ‘Income’ (IN), blue: “Majestic” (MJ), and growing system by fill: solid = soil, striped = hydroponic. Total xylem area was calculated by measuring the cross-sectional area of all individual xylem elements within each section, summing these values, and averaging per treatment. Cultivar-based differences were evident, while differences attributable to growing system were largely absent. Panel B: Average individual xylem diameter (µm) per treatment and root order. As with total xylem area, observed differences in individual xylem diameter reflected cultivar-based variation rather than growing system effects.

**Figure 4:**
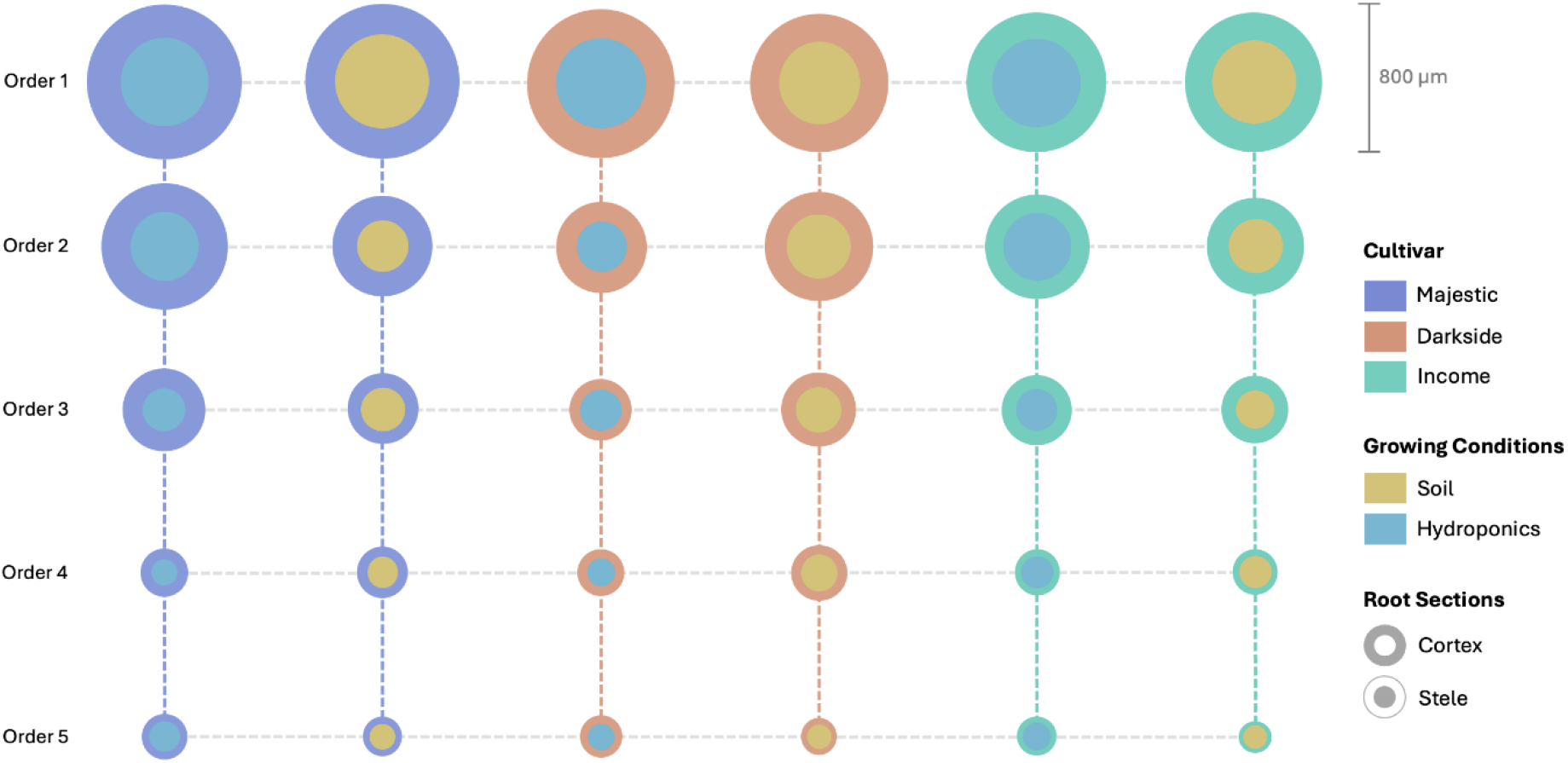
Root cross-sectional anatomy of spinach (*Spinacia oleracea*): stele and cortex diameter across root orders and growing systems. Bubble plot illustrating the relationship between stele diameter and cell (cortex) diameter across root orders, cultivars, and growing systems. Cultivars are denoted by the color of the outer circle, blue: ‘Majestic,’ orange: ‘Darkside,’ teal: ‘Income’, and growing system by the color of the inner circle: light blue = hydroponics, gold = soil. The inner circle represents the mean stele diameter (µm) per treatment (n = 4), and the outer circle represents the mean cell diameter (µm) of the root cross-section. Circle sizes are proportional to actual measurements; the scale bar represents 800 µm. Measurements were collected using a Zeiss Imager A2 microscope and processed in FIJI. Rows represent root order descending from first (top) to fifth (bottom); columns represent cultivar and growing system combinations. The consistent proportional relationship between stele and cell diameter across orders suggests that the vascular ratio may be influenced more by growing system than by cultivar.

#### 3.3 Environmental Conditions

Data collected of environmental conditions included temperature, light intensity, and dissolved oxygen levels. Average temperature was 25.2° C for the duration of the experiment. Light intensity of supplemental light averaged a range of 250-270 μmol/m^2^/s for the entire forty-five days. Dissolved oxygen concentration was measured in several sump tanks and reported at 5.7-5.9 mg/L.

## 4. Discussion

Root system architecture is a primary determinant of plant resource acquisition, and its modification in response to cultivation conditions has well-documented consequences for aboveground biomass production (Lombardi et al., 2021; Balliu et al., 2021). The consistent differences in fresh and dry leaf biomass observed between hydroponic and substrate-based systems in this study prompted an examination of belowground traits to identify which root system modifications contributed to these contrasts. Traits of interest included total root system length, root diameter, xylem anatomy, and root order distribution.

Morphological analysis revealed that hydroponic cultivation systems induced specific architectural shifts primarily characterized by longer, and finer root systems, whereas soil-grown plants were defined by thicker root systems dominated by shorter archetypes. Contrastingly, anatomical investigations indicated that internal root traits including stele dimensions and xylem diameter and area remained largely conserved across growth systems, displaying largely inconsistent, minor, variations among cultivars and root orders. The combination of such morphological plasticity and anatomical stability are consistent with the idea that fresh and dry leaf biomass differences between cultivation systems arose primarily from root system morphological adjustments rather than internal alterations.

### 4.1 Morphological root system differences are linked to cultivation systems

In our growth conditions, morphological distinctions in root systems among the cultivars ‘Income’, ‘Darkside’, and ‘Majestic’ were predominantly linked to differences in cultivation system: hydroponic versus soil, rather than by inherent cultivar-specific differences. Across all cultivars, hydroponically grown plants consistently exhibited finer, longer root systems with greater total dry root biomass and fresh-leaf weight relative to their soil-grown counterparts (Fig. 2A and Supplementary Fig. 2B).

Roots developed under hydroponic conditions displayed significantly greater (∼200% in harvest 2, ∼30% in harvest 3) total root length, dominated by higher order laterals (third to fifth order), which collectively work to expand the total absorptive surface area (Fig. 2D). This architecture closely resembles the “subsoil foraging ideotype,” characterized by extensive, finer, and exploratory roots optimized for efficient nutrient and water acquisition (Lombardi et al., 2021; Lynch, 2013). Although the SFI was originally defined in the context of field-grown crops, the present data suggest that functionally analogous architectural strategies emerge under NFT hydroponic conditions (Balliu et al., 2021; Liu et al., 2024). Recent literature indicates that such traits likely arise from the unique environmental conditions of hydroponic systems including high dissolved oxygen levels, low or no mechanical impedance, and readily available nutrients which initiate increased lateral root growth and elongation (Chen et al., 2022). Indeed, our DO measurements indicated a range of 5.7-5.9 mg/L, consistent with prior literature indications of DO in the range of 5.5-7.5 mg/L in hydroponics. The resulting proliferation of fine roots enhances nutrient uptake efficiency by increasing the potential root-nutrient solution interface, resulting in more advantageous absorption dynamics (Liu et al., 2024). In contrast, soil-grown roots exhibited a coarser morphology with thicker axes predominantly composed of lower order roots (first to third order). This configuration, though potentially beneficial for mechanical stability and storage, limits overall root length and exploration potential, thereby constraining nutrient and water uptake efficiency in soil media. Although some cultivar level variation was evident: ‘Income’ and ‘Darkside’ produced on average finer roots than ‘Majestic’ (0.35cm vs 0.45cm root diameter for harvest 2, 0.55cm vs 0.65cm for harvest 3), the environmental modulation of morphology clearly outweighed cultivar specific genetics (Fig. 2C). These morphological contrasts between cultivation systems were mirrored in aboveground traits, as hydroponically grown plants demonstrated substantially higher fresh leaf biomass across all cultivars.

These observations highlight the pronounced root morphological plasticity expressed by all three cultivars. That environmental conditions appear to exert a stronger influence on root architecture than cultivar background supports the notion of root systems as highly responsive, adaptive organs. Similar environmentally induced differences have been reported in lettuce (Lei & Engeseth, 2021), and tomato (Ayarna et al., 2020), where hydroponic cultivation promotes the proliferation of fine, highly branched roots, whereas soil cultivation promotes thicker, mechanically reinforced roots suited to sparser, heterogenous resource distribution (Sasse et al., 2020).

These mechanistic responses are well-documented in the literature. Hydroponic style conditions, especially constant hydration to the elongation zone or Root Apical Meristem (RAM) increases cell expansion and the feeding of the meristematic zone for cell division. The low impedance causes cells in both the RAM and root cap to be subject to reduced pressure and in response, they lignify less. Thus, the primary RAM shifts its growth style to one of sustained elongation over root thickening (Que et al., 2018). This is consistent with the reduced root diameter and greater total root length observed in hydroponic plants (Fig. 2C, 2D), and with the greater fresh leaf biomass relative to dry root biomass observed across cultivars (Figs. 2A and Supplementary Fig. 2B).

This capacity for structural adjustment suggests that root system plasticity is a strong determinant of plant performance under contrasting cultivation systems. An interesting further avenue of exploration along these lines is whether plants grown in soil can serve as a proxy or point of reference for the behavior of that same plant in a hydroponic style system.

### 4.2 Alterations of vascular and root anatomy are minor when compared to morphological differences in response to growth system

Anatomical characteristics of the root system revealed a complex interaction between developmental order, cultivar, and cultivation system. Several variables of interest including total xylem area (p<.002) and vascular ratio (p<.03) had significant two-way interactions between cultivar and root order and cultivar and system respectively. However, when growth system was added to the former model, and order to the latter model, creating three-way interactions, they became insignificant. It becomes clear then that while some anatomical plasticity is present, it is linked to cultivar more than growth system. Furthermore, these internal structural differences did not translate into proportional changes in fresh and dry leaf biomass. Whereas morphological analyses demonstrated that hydroponic cultivation was highly linked to the promotion of finer, exploratory (development of higher root orders) root systems, the anatomical data suggest that vascular organization remains comparatively stable and less responsive to environmental modulation (Balliu et al., 2021; Li et al., 2018).

A significant three-way interaction among root order, cultivar, and system were detected for xylem diameter (p<.003), average individual xylem area (p<.006) and near significance for stele diameter (p<.06). reflecting a modest degree of anatomical plasticity. However, the trend and magnitude of these differences were inconsistent across orders and cultivars, indicating that while anatomical traits respond to environmental conditions to some degree, such responses were highly context dependent (Figs. 3 and 4). For instance, hydroponically grown roots often displayed slightly larger stele diameters (475 μm Hydroponic vs 400μm Soil Order 1) and higher vascular tissue ratios (55% Hydroponic vs 50% Soil Order 1, particularly in first and second-order roots, consistent with an early developmental enhancement of xylem conductive capacity (Balliu et al., 2021). Yet, this trend did not persist uniformly across higher root orders, where soil-grown roots occasionally exhibited comparable or greater vascular dimensions (Stele Diameter 3^rd^,4^th^, 5^th^ orders (Figs. 4 and 5).

**Figure 5:**
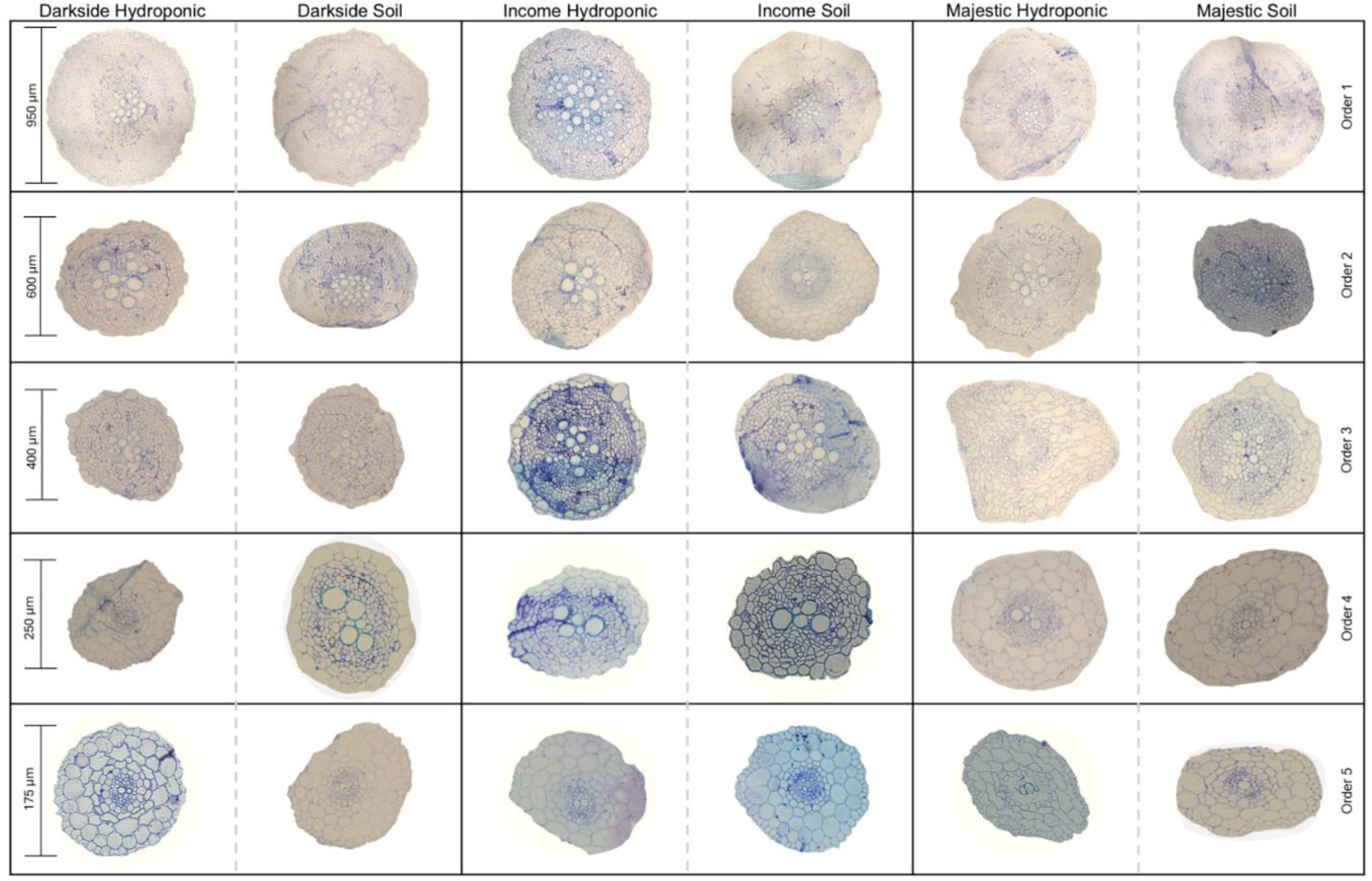
Representative root cross-sections of spinach (*Spinacia oleracea*) across root orders and growing systems. Light microscopy images of toluidine blue-stained root cross-sections for all three cultivars, both growing systems, and all five root orders. Columns represent cultivar and growing system combinations: ‘Darkside’ Hydroponic, ‘Darkside’ Soil, ‘Income’ Hydroponic, ‘Income’ Soil, ‘Majestic’ Hydroponic, and ‘Majestic’ Soil. Rows represent root orders 1 through 5 (top to bottom). Scale bars on the left indicate the approximate diameter at each root order: 950 µm (Order 1), 600 µm (Order 2), 400 µm (Order 3), 250 µm (Order 4), and 175 µm (Order 5), reflecting the progressive decrease in root diameter with increasing order. Sections were prepared by fixation, dehydration, and embedding in JB4 resin, followed by sectioning at 1.5 µm thickness and staining with toluidine blue. Imaging was performed using a Zeiss Imager A2 microscope. The large circular structures visible within the stele represent xylem vessels, which are notably larger and more prominent in lower-order roots. Staining intensity and cellular organization vary across cultivars and orders, consistent with the quantitative anatomical data presented in Figures 3 and 4.

The limited consistency in anatomical adjustment stands in stark contrast with the clear and systematic morphological divergence observed between systems. The proliferation of higher order fine roots under hydroponic cultivation appears to enhance resource uptake primarily through surface area expansion rather than through basic modifications to vascular structure (Li et al., 2018). In this context, the modest increases in xylem and stele dimensions in hydroponic roots likely serve to support, rather than drive, enhanced shoot biomass production (Figs. 3 and 4) (de Moraes et al., 2022; Duan et al., 2020; Que et al., 2018). The greater fresh weight of hydroponically grown plants therefore appears to arise principally from the architectural and physiological efficiencies including increased absorptive interface and reduced mechanical impedance of hydroponic media rather than from intrinsic differences in vascular anatomy (Balliu et al., 2021; Awika et al., 2021).

Comparable trends have been documented across several vegetable crop species where environmental conditions induce morphological but not proportional anatomical plasticity. In one study of lettuce, for example, hydroponic cultivation promotes extensive fine root proliferation with only modest changes in xylem vessel size or density (Lei & Engeseth, 2021). However, in other crops such as wheat, the opposite has occurred, wherein the xylem of plants grown in water-abundant environments has been wider, with increased total areas (Bresta et al., 2011). This has been hypothesized to be due to the relationship between radius and flow through the capillary tubes in relation to xylem safety vs efficiency. It has also been suggested that there is a relationship between xylem conductance derived from xylem anatomy that is altered in response to environmental conditions such as drought (Hernandez-Espinoza & Barrios-Masias, 2020). It would thereby be an acclimation response which could play a role between water transport and gas exchange in order to maximize drought survival (Bresta et al., 2011). In tomato, this trend became clear, with total vascular area significantly higher in hydroponic crops than in soil (MacLellan, 2019). There is a considerable gap of research in the comparison of growth systems with regard to vegetable crops (Dsouza et al., 2023; Wu et al., 2022). To better understand these mechanisms, including whether they are decoupled and, if so, in which crop classes, more studies are needed. Examples include lignation vs expansion, signaling processes, exudate applications, and environmental stress conditions altering root architecture (Dsouza et al., 2023). In this study, the parallels with the experiments formerly described above reinforce the interpretation that in spinach, the system-induced gains in vegetative biomass are primarily morphological in origin. As such, the cultivars in this study express a conserved anatomical framework capable of supporting substantial morphological flexibility across various cultivation systems.

Collectively, these results illustrate that while spinach root morphology exhibits clear environmental plasticity, the underlying anatomical framework remains comparatively conserved (Fig. 5). In our limited study, vascular organization in all three cultivars adapts subtly to cultivation systems but does not appear to be the primary determinant of overall plant productivity. Instead, the coordination between morphological plasticity and stable internal anatomy underscores an integrated developmental strategy. Such a strategy enables root systems to modify their external development for environmental acquisition demands, while maintaining sufficient internal structure (vascular tissue) to sustain transport needs and structural integrity. Such a decoupling between external morphology and internal anatomy may represent an adaptive balance mechanism between flexibility and functionality, possibly ensuring efficient resource use without compromising physiological stability across cultivation systems. However, this may not be the case in all scenarios or environments. The small sample size for microscopic analysis, along with the loss of some of the 3-5^th^ order roots in the cleaning and scanning process may lead to differing findings under other growth conditions or cultivars. Future studies with larger sample sizes and varied growth conditions should seek to reinforce these findings.

### 4.3 Advancing toward a fine-root dominant ideotype

The development of the Subsoil Foraging Ideotype (SFI) in spinach, most clearly expressed in this study in the cultivar ‘Income’, illustrates a root architecture optimized for efficient environmental exploration and growth across all five orders. Characterized by a dominant, vertically penetrating primary root accompanied by a dense proliferation of fine lateral roots, such a morphology promotes extensive root exploration and improved access to subsoil water and nutrients (Koevoets et al., 2016; Lynch, 2013). Such an archetype reflects ideotypes reported in other species under resource-scarce conditions, wherein an initial development of first and second order roots gives way to fine root proliferation to enhance resilience to transient drought and nutrient depletion (Lombardi et al 2021; Lynch 2013).

In CEA systems, these traits carry distinct advantages across both high and low-technology production systems. Within contexts of high-tech hydroponic or vertical farming, the exploratory foraging tendency of ‘Income’s root system may be simulated through root-zone management or substrate design that mimics subsoil stratification. Enhanced root exploration under such conditions may contribute to the elevated nutrient uptake efficiency and rapid biomass accumulation characteristic of ‘Income’ relative to other cultivars (Balliu et al., 2021). This is visible in tracking of dry root weights compared to fresh leaf weight along periodic harvest. Cultivar ‘Income’ had the highest dry root weight in the 2^nd^ and 3^rd^ harvests, ∼3g and ∼7g, respectively. In contrast, the next highest cultivar was 2.5g and 4.5g, respectively. When compared to the highest fresh leaf weight for the 2^nd^ and third harvest, ‘Income’ was again the highest at 40g and ∼80g vs 30g and 55g, respectively, for the next highest cultivar. It thus becomes clear that there is an impact on the proliferation of the fine roots on the fresh leaf growth, which is the aim of CEA. This accelerated growth may enable high-tech systems to achieve a greater number of growth cycles per year, translating to improved turnover rates and production scalability. Consequently, the SFI’s inherent capacity for vigorous, fine-root proliferation aligns closely with the performance demands of intensive, year-round cultivation frameworks, such as those in urban agriculture settings (Lynch, 2022).

In contrast, under lower-tech or soil-based CEA systems, the same SFI traits may offer adaptive advantages under fluctuating environmental conditions. The ability to penetrate deeper soil layers allows plants to access hidden pools of moisture and nutrients beyond the reach of shallower-rooted phenotypes, buffering against drought stress and nutrient variability intensified by climatic instability (Kou et al., 2022, Lynch, 2022). The same growth patterns as described in the above paragraph detail the ability for these types of cultivars to achieve significant fresh leaf biomass as compared to other cultivars. Based on prior literature, this suggests enhanced foraging efficiency, which may support sustained productivity in resource-limited settings, thereby reinforcing food system resilience at smaller operational scales (Benitez-Alfonso et al., 2023).

Across both growth system types, SFI represents a morphological and functional convergence of efficiency and adaptation. The fine-root dominance and growth architecture it possesses not only underpin robust nutrient acquisition strategies but also illustrate a broader capacity for plastic responses to the cultivation environment. The ‘Income’ cultivar’s manifestation of this ideotype positions it as a candidate for targeted breeding toward climate-resilient and systemically adaptable spinach varieties. Further investigation into the physiological and anatomical foundations of SFI function, particularly its interactions with substrates and nutrient uptake, will be key to optimizing its capabilities in controlled-environment systems (Sharma et al., 2024). Furthermore, explorations of internal anatomy, especially in xylem and phloem development could positively influence breeding and development. Additional avenues of exploration include coupling growth-system microbiota with root exudates to further understand their impact on root system morphology (Camli-Saunders & Villouta, 2025). Ultimately, the integration of root system architecture in agriculture decision-making may enable both high and low-tech CEA systems to enhance their productivity while maintaining resilience under increasing climatic pressures.

## Supporting information

Supplemental Figures + Protocol

## CRediT Author Contributions

*Deniz Camli-Saunders*: Conceptualization; Methodology; Investigation; Data curation; Formal analysis; Visualization; Writing – original draft.

*Ava K. Russell*: Investigation; Data curation.

*Camilo Villouta:* Conceptualization; Methodology; Funding acquisition; Resources; Supervision; Writing – review and editing.

## Data Availability

The raw data supporting the conclusions of this article are openly available in the Zenodo repository at the following DOI: 10.5281/zenodo.19224067

## Declaration of Generative AI and AI-Assisted Technologies in the Manuscript Preparation Process

During the preparation of this work, no generative AI and AI-assisted technologies were used.

## Acknowledgements

The authors would like to thank the Friedman Lab at the Arnold Arboretum, Harvard University, for their assistance with JB-4 plastic resin sectioning training and troubleshooting.

