## Supplemental Figures + Protocol for "Comparative analysis of root morphology in several spinach (Spinacia oleracea) varieties: Field vs Hydroponic growth systems"

**Supplementary Figure 1 Title:** Root architectural complexity metrics of spinach (*Spinacia oleracea*) across harvests and growing systems

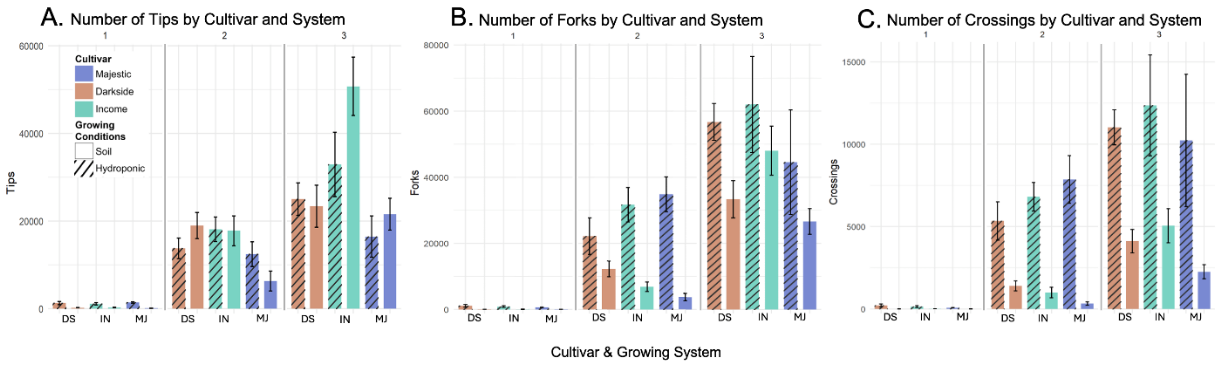


Supplementary Figure 1.

**Supplementary Figure 1 Caption:** Bar plots of root architectural complexity metrics across three harvests, cultivars, and growing systems. Cultivars are denoted by color: blue: 'Majestic' (MJ), orange: 'Darkside' (DS), teal: 'Income' (IN), and growing system by fill: solid = soil, striped = hydroponic. Harvest 1 (day 15), 2 (day 30), and 3 (day 45) are shown left to right within each panel. Error bars represent ± 1 standard error of the mean (n = 10). Metrics were collected using the WinRhizo root scanner. Panel A: Total number of root tips. Panel B: Total number of root forks. Panel C: Total number of root crossings. All three metrics increased substantially with harvest, reflecting overall root system growth over time. 'Income' showed consistently higher values across metrics by harvest 3, particularly under hydroponic conditions.

**Supplementary Figure 2 Title:** Additional root biomass and morphological measurements of spinach (*Spinacia oleracea)* across harvests and growing systems

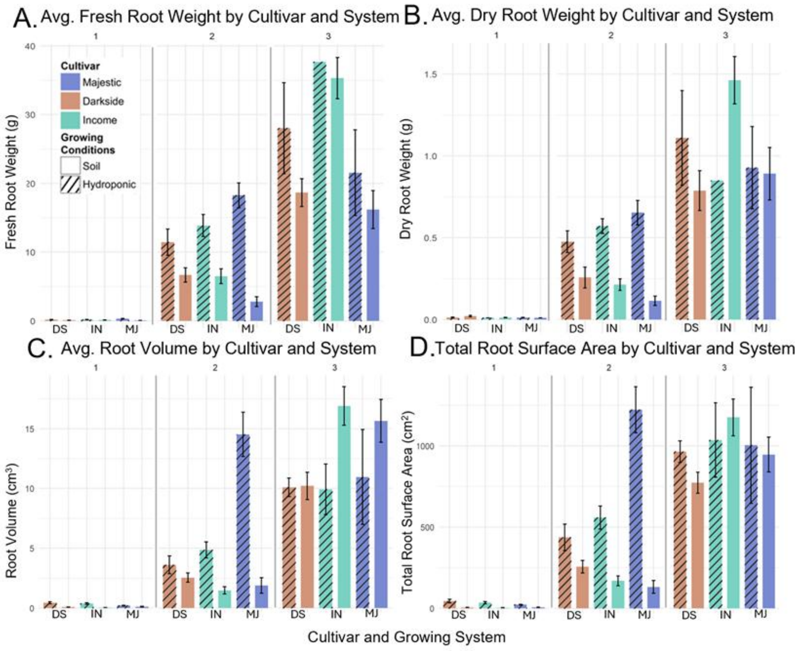


Supplementary Figure 2.

**Supplementary Figure 2 Caption:** Bar plots of additional root biomass and morphology metrics across three harvests, cultivars, and growing systems. Cultivars are denoted by color: blue: 'Majestic' (MJ), orange: 'Darkside' (DS), teal: 'Income' (IN), and growing system by fill: solid = soil, striped = hydroponic. Harvest 1 (day 15), 2 (day 30), and 3 (day 45) are shown left to right within each panel. Error bars represent ± 1 standard error of the mean (n = 10). Root metrics were collected using the WinRhizo root scanner. Panel A: Average fresh root weight (g) measured immediately after harvest. Panel B: Average dry root weight (g) after 24 hours at 60°C. Panel C: Average root volume (cm³). Panel D: Total root surface area (cm²). All metrics increased with harvest, consistent with overall root system growth over time. Hydroponically grown plants showed notably higher root volume and surface area by harvest 3, particularly in the 'Majestic' and 'Income' cultivars, suggesting greater root system development under hydroponic conditions.

**Supplementary Figure 3 Title:** Xylem number and individual xylem area of spinach (*Spinacia oleracea*) roots across root orders and growing systems

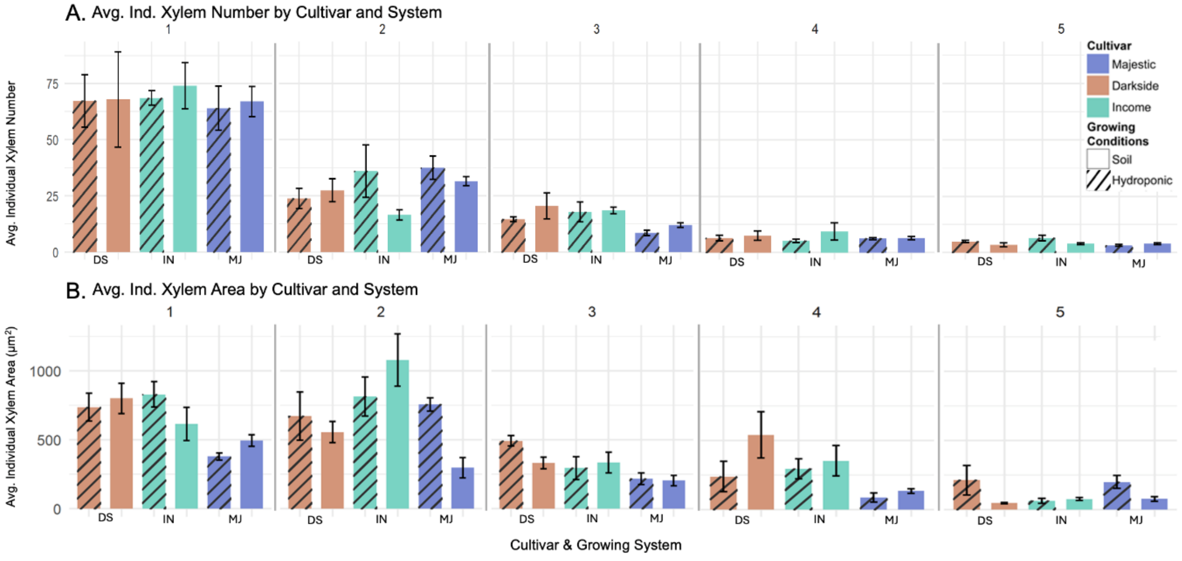


Supplementary Figure 3.

**Supplementary Figure 3 Caption:** Bar plots of individual xylem number and area across root orders, cultivars, and growing systems. Cultivars are denoted by color: blue: 'Majestic' (MJ), orange: 'Darkside' (DS), teal: 'Income' (IN), and growing system by fill: solid = soil, striped = hydroponic. Root orders 1 through 5 are shown left to right within each panel (n = 4). Error bars represent ± 1 standard error of the mean. Sections were prepared and imaged as described in Figure 3. Panel A: Average number of individual xylem elements per root cross-section. Xylem number decreased progressively with increasing root order across all cultivars and systems, consistent with the reduced vascular complexity of higher-order roots. Panel B: Average individual xylem element area (µm²). A similar decreasing trend was observed with increasing root order. Both panels suggest that differences in xylem number and area are primarily driven by root order rather than by cultivar or growing system.

**<u>JB-4 Dehydration, Infiltration, Embedding Protocol</u>**

Adapted from Sigma-Aldrich Technical Bulletin for JB-4

*All steps should be conducted under a fume hood and with gloves*

**Fixation/Dehydration/Infiltration**

1. Samples should be fixed in 4% Paraformaldehyde solution for 24 hours in 4°C refrigerator on a shaker. Use 5 psi vacuum for period of 5 minutes to aid in infiltration (x3)
2. After 24 hours, remove from refrigerator and wash 3x with PBS. Washes should be done once per hour with sample tubes on shaker plate
3. After washes, dehydrate samples up to 70% ethanol with the following steps: 10% EtOH, 20%, 30%, 50%,70%- at least one hour per step
4. Can stop here for long term storage
5. Continue to dehydrate to 85%, 95% x3, one hour per step again, continue shaking
6. Bring samples to 100% MonoA solution through the following steps: 3:1 (95% EtOH:MonoA), 1:1 (95% EtOH:MonoA), 1:3 (95% EtOH:MonoA), 100% MonoA. Each step should take 24 hours in length, continue shaking.
7. Change out 100% MonoA solution 1x per week (x3)

**Embedding**

1. Prepare 100 mL of JB4 infiltration solution using the following: 100 mL solution A (monomer) and 1.25g Benzoyl Peroxide Catalyst using a magnetic stirrer until fully dissolved.
2. To prepare embedding solution, use 25 mL of fresh infiltration solution mixed with 1.0 mL of JB4 solution B and embed immediately as catalyzation is rapid.
3. Place root sample in block an add embedding solution, can use toothpick to arrange sample as desired
4. It is key to exclude oxygen while samples polymerize. Use nitrogen gas or light vacuum (10-15psi) to create anoxic environment for ideal polymerization. Samples polymerize fully in 45 mins to an hour. Large samples may take longer.
5. Polymerized samples may have thin liquid film on top, place block in sealed container with desiccant beads to remove.
6. Polymerized samples can be glued to stubs for sectioning.
